# The *Drosophila* seminal Sex Peptide can associate with rival as well as own sperm and provide function for SP in polyandrous females

**DOI:** 10.1101/2020.02.20.958108

**Authors:** Snigdha Misra, Mariana F. Wolfner

## Abstract

In populations in which females tend to mate with more than one male, sperm competition and cryptic female choice can occur, triggering biases in sperm use and influencing males’ paternity share outcome of the mating. This competition occurs in the context of molecules and cells of male and female working interdependently towards the common goal of optimal fertilization. For example, a male’s seminal fluid molecules modify the female’s physiology to increase reproductive success. However, since some of these modifications induce long-term changes in female physiology, this can indirectly benefit rival males. Indeed, rival males can tailor their ejaculates accordingly, minimizing the energy cost of mating. Here we investigate the direct benefits that seminal fluid proteins from an ejaculate of one male can confer to sperm of a rival. We report that Sex Peptide (SP) that a female receives from one male can bind to sperm from a prior mate, that were already stored in the female. Moreover, the second male’s SP can restore fertility and facilitate efficient sperm release or utilization of sperm received from the first male that had been stored in the female. Thus, SP from one male can directly benefit another and as such is a key molecular component in the process of inter-ejaculate interaction.

## Introduction

In many animal species, females mate with more than one male. This polyandry lays the foundation for sperm competition, in which ejaculates from rival males compete for fertilization opportunities [1,2]. These conflicts and associated cryptic female choice can drive the evolution of male traits including optimal sperm numbers, morphology, and seminal protein sequences [3–5].

Against the backdrop of these conflicts, male and female molecules and/or cells must also work together to ensure reproductive success. How efficiently sperm interact with the egg and instigate successful fertilization or embryo support (where relevant) is key to successful fertility. Accordingly, males have evolved molecular mechanisms to trigger physiological changes in females that increase the reproductive success of the mating pair. Seminal fluid proteins (SFPs) are crucial regulators of these changes. SFPs are produced within glandular tissues in the male reproductive tract and are transferred to females along with sperm during mating [6–11]. Within a mated female, SFPs mediate an array of post-mating responses such as, in insects, changes in egg production, elevated feeding rates, higher activity or reduced sleep levels, long-term memory, activation of the immune system and reduced sexual receptivity [12–18].

The ability of a male’s SFPs to induce long-term changes in the mated female enhances that male’s reproductive success. For example, the seminal Sex Peptide (SP) of male *Drosophila* binds to his sperm stored in the female, persisting there for approximately ten days [19]. This binding of SP to sperm is aided by the action of a network of other SFPs-“LTR-SFPs” [10,20,21]. The active region of SP is then gradually cleaved from sperm in storage, dosing the females to maintain high rates of egg laying, decreased receptivity to remating [19], increased food intake, and slower intestinal transit of the digested food to facilitate maximum absorption and production of concentrated faeces [12,22–24]. However, induction of these changes can also indirectly benefit his rival, as the female’s physiology will have already been primed for reproduction by her first mate’s SFPs. Such indirect benefits to the second male have been suggested to explain the tailoring of the ejaculate by males that mate with previously mated females [11,25–27]. For example, the *Drosophila* seminal protein ovulin increases the number of synapses that the female’s Tdc2 (octopaminergic) neurons make on the musculature of the oviduct above the amount seen in unmated females [28]. This is thought to sustain high octopaminergic (OA) signalling on the oviduct musculature of mated female, allowing increased ovulation to persist in mated female, even after ovulin is no longer detectable in the female. Therefore, males mating with previously mated females need transfer less ovulin than males mated to virgin females, presumably because it may be less necessary, as they benefit from the ovulation stimulating effect of ovulin from the prior mating. In another example, prior receipt of Acp36DE can rescue sperm storage of a male that lacks this SFP [29,30].

The benefits to the second male described above are indirect consequences of the first male’s SFPs effect on female’s physiology. The second male is thus the lucky beneficiary of the first male’s SFPs effect on the female. However, it is unknown whether a male could directly benefit from a rival’s SFPs, for example, whether the latter could associate with and improve the success of another male’s sperm. There was some suggestion that this might occur from the phenomenon of “copulation complementation”[31] in which a female *Drosophila* singly-mated to a male lacking SFPs did not produce progeny unless she was remated to a male who provided SFPs. This suggested that something from the second mating allowed the first male’s sperm to be used. However, the molecular basis for this phenomenon was unknown. Also suggestive that a male could benefit from a rival’s SFPs, Avila et al [6] reported that *SP*-null males were better at defensive sperm competition than controls: mates of SP null males sired significantly more progeny (higher P1) than mates of control males when females were mated with the *SP-*null, or control male, and then with a wild type (wt) competitor. The higher P1 of *SP*-null males in this situation occurred because their mates had retained more of their sperm than of control males, due to the requirement for SP for release of sperm from storage, and then the release and use of those sperm when the females received ejaculate from the second, wt male.

Here, we report that SP, a *Drosophila* SFP received from a second male can bind to a prior male’s SP-deficient sperm and restore his fertility, including sperm release from storage and changes in the female’s behavior. Our results reveal direct benefits that previously stored sperm from the first (or prior) male can receive from the second (or last) male’s ejaculate during the course of successive matings. Our results also establish SP as a crucial long-term molecule that facilitates this inter-ejaculate interaction, and SP-sperm binding as the molecular mechanism that underlies copulation complementation in *Drosophila*. We discuss these findings in relation to sperm competition and the possibility of copulation complementation in nature.

## Results

### 1. Sex peptide from one male can associate with sperm from another

In matings with wt males, SP binds to sperm that it enters the female with. However, we wondered if sperm stored by mates of *SP*-null males, that lack bound SP, could become decorated with SP from a second male even if he did not provide sperm. If so, this would mean that SP from a second (spermless) male can bind to sperm from a prior male, already present in female tract (Fig 1. Cartoon).

**Figure 1.**
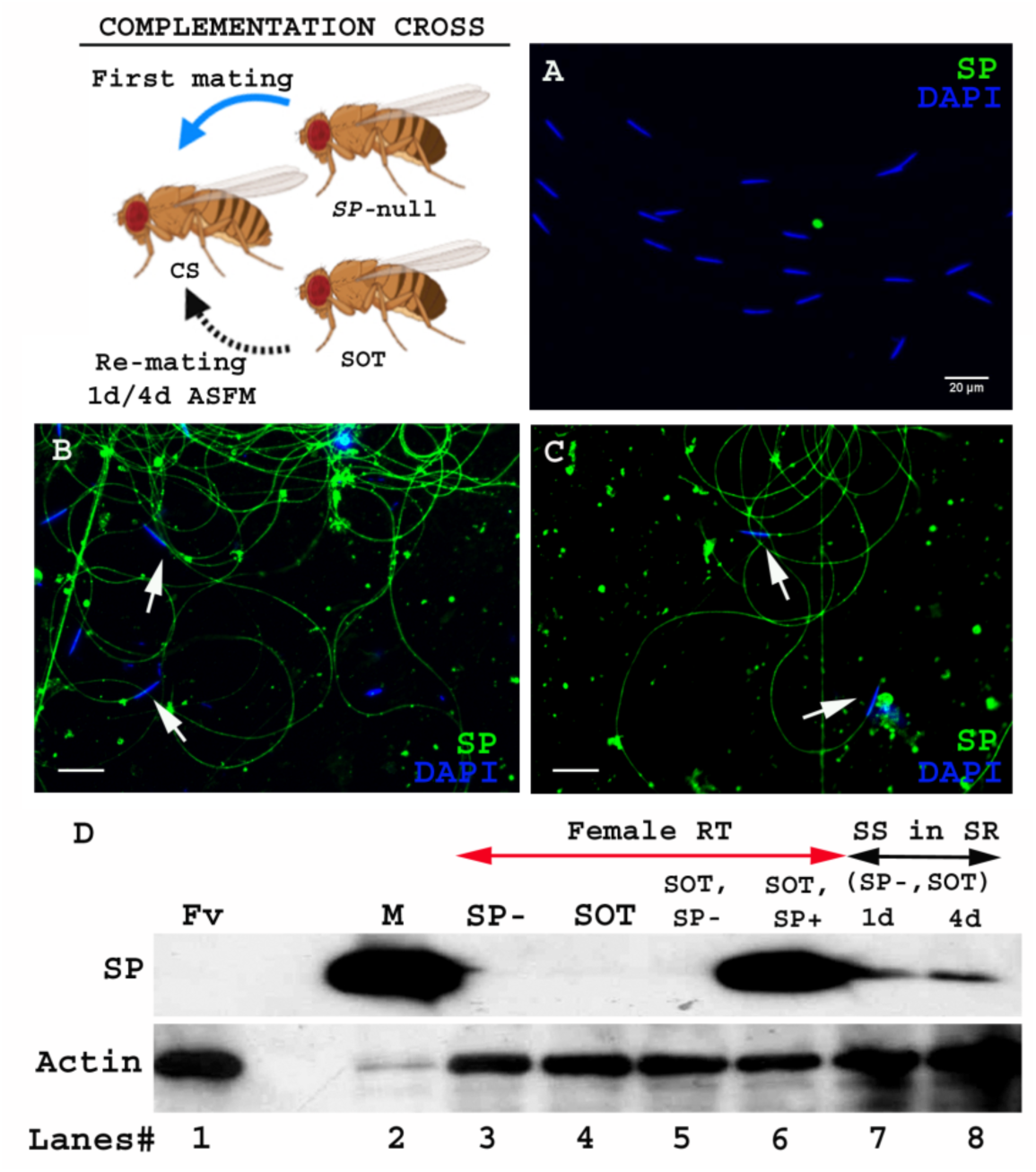
SP from a second male can bind to SP deficient sperm of previous male stored within a mated female. **Cartoon**: Pictorial representation of the crossing scheme (fly images from Biorender). Wild type (CS) females were first mated to an *SP*-null male and then, at the indicated time, to a spermless (*sot*) male. Sperm heads were stained with DAPI (blue) and SP visualized with Alexa fluor 488, staining the sperm tail (green) and sperm head (cyan; overlapping blue/green). **(A)** Sperm from females singly mated to *SP*-null males, 1d ASM. **(B)** Sperm from females mated to *SP*-null males, remated to spermless males at 1d ASFM and **(C)** at 4d ASFM, both frozen 2h ASSM. White arrows indicate sperm heads. Bar=20µm **(D)** Western blot lanes# **1:** Fv, reproductive tract (RT) of virgin female (negative control; n=5), **2:** M, a pair of male accessory gland (positive control; n=1), **3:** SP-, reproductive tracts of females mated to *SP*-null males, 2h ASM (n=5), **4:** SOT, reproductive tracts of females mated to spermless males, 1d ASM (n=5), **5:** SOT, SP-, reproductive tract of females mated to spermless males and then remated to *SP-*null males, 1d ASFM (n=8 RT), **6:** SOT, SP+, reproductive tract of females mated to spermless males and then remated to control (SP+) males at 1d ASFM, frozen 2h ASSM (positive control; n=8 RT), **7:** (SP-, SOT), 1d and **8:** (SP-, SOT), 4d sperm isolated from the seminal receptacle of females mated to *SP*-null males and then remated to spermless males at 1d ASFM and 4d ASFM, frozen 2h ASSM (n=15 SS). Actin served as loading control.

To test whether SP from a second male can bind to SP-deficient sperm stored by mates of *SP*-null males, we first confirmed that no SP was detectable on sperm stored in females that had singly-mated to *SP-*null males (Fig 1A). We then examined whether SP was detected on such sperm if the female subsequently remated to a spermless male (who provided SP). We observed SP bound to the stored sperm following such rematings at either 1d (Fig 1B) or 4d (Fig 1C) after the original SP-less mating. We confirmed these findings with western blotting. Sperm stored in seminal receptacles of females that had mated to *SP*-null males and subsequently remated to spermless males were dissected and probed for the presence of SP. Consistent with our immunostaining data, SP was detected in samples of *SP-*null male’s sperm from females that had remated to spermless males at 1d or 4d after the start of first mating (ASFM; Fig 1D, lanes 7 and 8). Thus, SP from a second male can bind to SP-deficient sperm stored from a prior male. To see if the mating order was important, we carried out the reciprocal cross, i.e. testing if SP deposited by a first male (spermless, in this scheme) could bind to sperm that were subsequently introduced by a second (*SP*-null) male (Fig 2. Cartoon). Spermless males transfer SP to the female tract after mating [32], but we did not detect any SP in females mated to spermless males by 1d after the start of mating (ASM; Fig 1D. lane 4). We saw no SP signal in sperm samples isolated from females that had mated to spermless males, and then subsequently to *SP-*null males at 1d ASFM (Fig 1D. lane 5). Our immunofluorescence data were consistent with our western blots: we saw no SP-sperm binding in females that mated first with a spermless male and a day later with *SP*-null male (Fig 2B). Therefore, if SP entered the female without sperm, it was unavailable to bind to sperm from a subsequent SP-deficient male.

**Figure 2.**
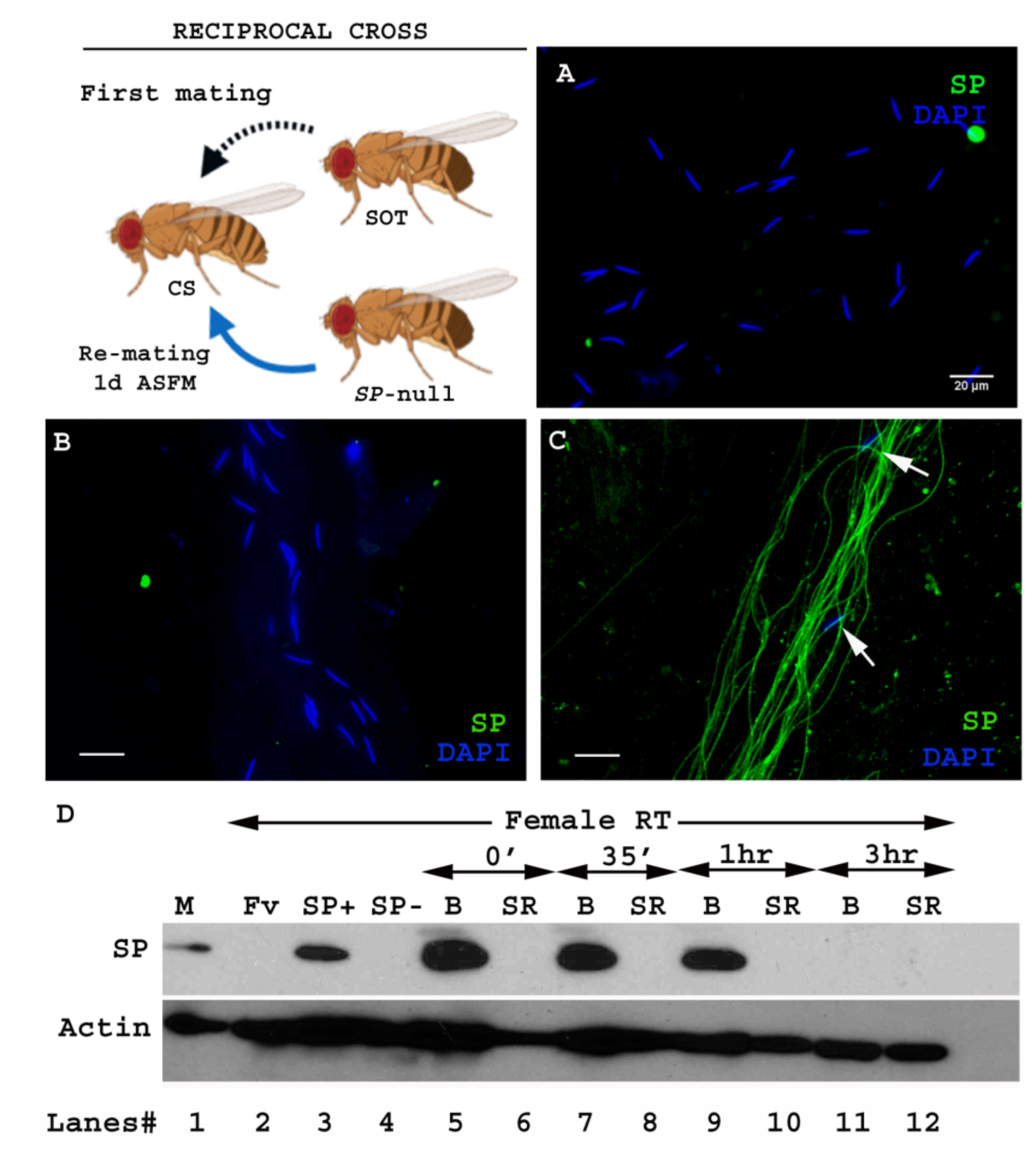
Sperm from second male are not bound to SP from a prior spermless male. **(Cartoon):** Pictorial representation of cross (fly images from Biorender), that is reciprocal of that in Fig 1. Females mated first with spermless (*sot*) male and then a day later with *SP*-null male that provided sperm. Sperm heads were stained with DAPI (blue) and SP visualized with Alexa fluor 488, staining the sperm tail (green) and sperm head (cyan; overlapping blue/green). **(A)** Sperm from females singly mated to *SP*-null males, 2hr ASM. **(B)** Sperm from females mated to spermless males and then remated to *SP*-null males, 1d ASFM. **(C)** Sperm from females mated to spermless males and then remated to SP+ males, 1d ASFM, serve as positive controls. Flies were frozen 2h ASSM. White arrows indicate sperm heads. Bar=20µm **(D)** Western blot lanes# **1:** M, a pair of male accessory gland (positive control; n=1), **2:** Fv, reproductive tract (RT) of virgin female (negative control; n=5), **3:** SP+, reproductive tract of females mated to control males (TM3 siblings of *SP*-null males; n=5; positive control), **4:** SP-, reproductive tract of females mated to *SP*-null males (n=5; negative control). **5-12:** Proteins from Bursa (B) or seminal receptacle (SR) from females mated to spermless males frozen at 0’(min) immediately after mating, 35’(min), 1hr, and 3hr ASM, respectively (n=15). Actin served as loading control.

We hypothesized that we did not see SP bound to sperm in this second (reciprocal) crossing scheme because by the time of the second mating SP from the spermless male was no longer present in the female at 1d ASFM, since it could not be retained without binding to sperm [19] and no sperm were being supplied by these first males. To circumvent this, we attempted to remate females that had previously mated to spermless males as soon as 3-6hrs ASFM. However, few females remated, likely due to the recent experience of copulation, or to the effects of pheromones from the previous mating [33,34]. In the few females that did remate, no SP-sperm binding was observed (Fig S1). Since the simplest explanation for these results was that SP transferred without sperm had disappeared from females by the time of the second mating, we performed western blotting to determine how long SP persists in the reproductive tract of females in absence of sperm. Females mated to spermless males were flash frozen at 0’(min) immediately after mating, 35’(min), 1hr, and 3hr ASM and their bursa (B) and seminal receptacle (SR) were dissected and probed for the presence of SP. We detected SP in the bursa protein samples at 0’(min) after mating, 35’(min), and 1hr ASM. (Fig 2D. lanes 5, 7, 9). However, SP was undetected in bursa or seminal receptacles of females at 3hr ASM (Fig 2D. lane 11, 12). Thus, we could not determine whether SP from mating with a spermless male could bind a second male’s sperm, because SP received from the first mating was lost from the female reproductive tract before a second mating could occur. Xue and Noll [31] reported that a similar cross (females mated first to spermless males and then to *Prd* males) also gave no progeny (showed no copulation complementation) which they proposed to be due to inactivation or early loss of SFPs in the absence of sperm. Our results, showing that SP can bind to stored sperm from a prior male, provide the molecular explanation for their observation.

### 2. SP from a second male restores fertility, inhibits receptivity and regulates optimal release of the first male’s sperm from storage

SP is needed for efficient sperm release and utilization from the female sperm storage organs [6]. We tested whether SP from a second male could restore the use of a first male’s sperm. Females mated to spermless males have no progeny (Fig 3A). Females singly-mated to *SP*-null males have significantly reduced numbers of progeny (Fig 3A. SE of diff = 8.043; p***=<0.001) relative to females mated to control males (Fig 3A), likely because lack of SP prevents increase in egg production [17,35,36] and release of sperm from storage [6]. However, females mated to *SP*-null males and then remated to spermless males at 1d (Fig 3B; p=0.2487) and 4d (Fig 3C; p=0.8618) ASFM had progeny levels similar to those of females that had mated to control (SP^+^) males and were subsequently remated to spermless males at the same time points. Thus, SP from a second (SOT) male could rescue the fertility defects that resulted from the lack of SP from an *SP*-null first male.

**Figure 3.**
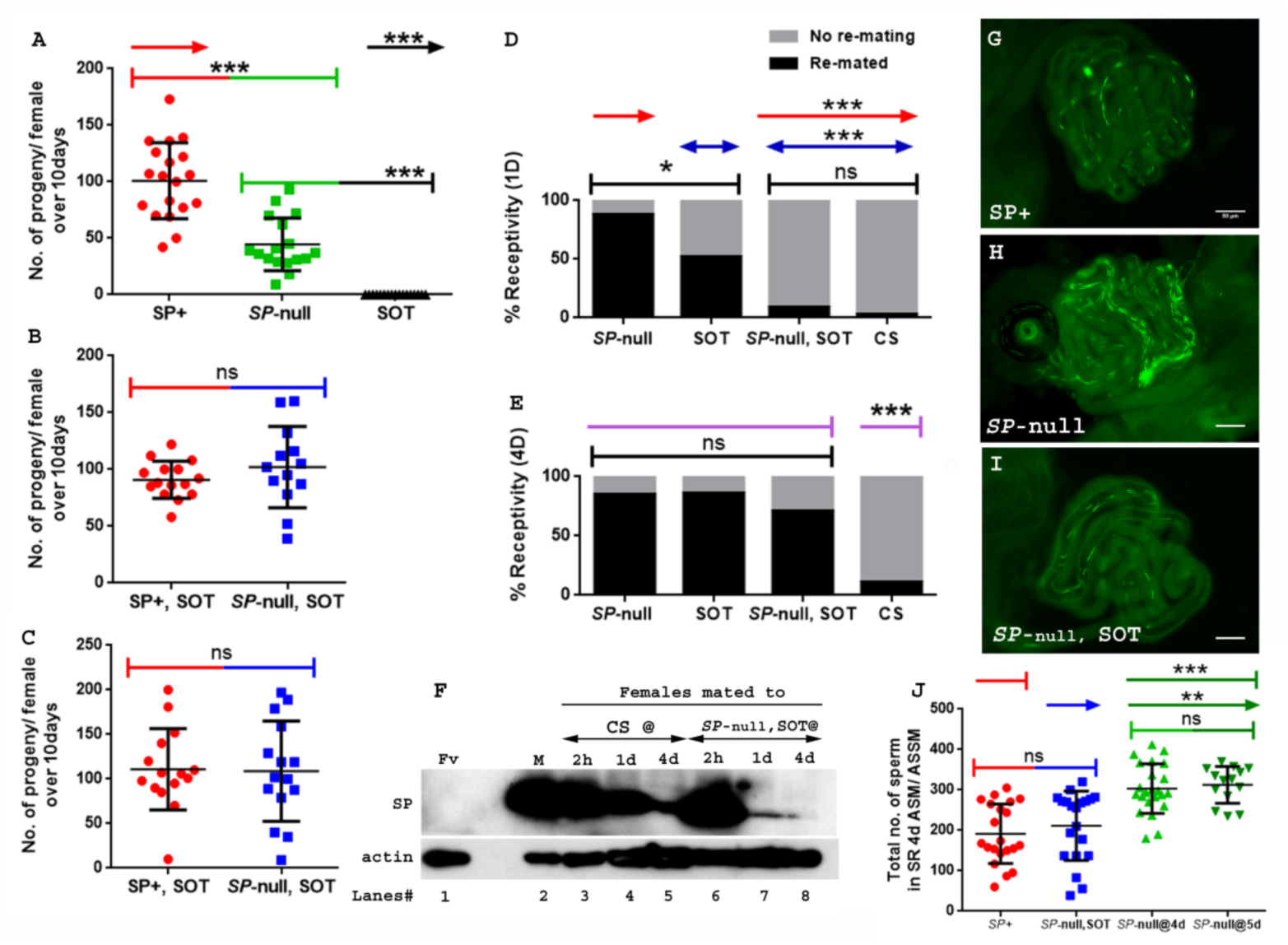
Remating with spermless males restores fertility, delays receptivity and optimizes efficient sperm release in females that previously mated to SP-null males. **(A)** Graphical representation of numbers of progeny produced by each female over the span of ten days, following mating to control (TM3 siblings of *SP-*null males: SP+; red), *SP*-null males (*SP*-null; green), or spermless males (SOT), p***=<0.001; n=15-20. **(B)** Fertility of females mated to *SP-*null males and then remated to spermless males at 1d ASFM (*SP*-null, SOT; blue, n=15-20) and **(C)** Fertility of females mated to *SP*-null males and then remated to spermless males at 4d ASFM (*SP*-null, SOT; blue, n=15-20) compared to females mated to control males and then remated to spermless males (SP+, SOT, red, ns=non significant). **(D)** Percentage receptivity of females mated to *SP-*null males and then remated to spermless males (*SP*-null, SOT) at 1d ASFM, when compared to females singly mated to *SP*-null males (red arrows), spermless (SOT, blue arrows) or CS males, 1d ASM (p*=<0.05; p***=<0.001; n=15-20). **(E)** Percentage receptivity of females mated to *SP*-null males and then remated to spermless males (*SP*-null, SOT) at 4d ASFM, when compared to females singly mated to *SP*-null males, spermless (SOT) or CS males (purple arrows), 4d ASM (p***=<0.001; n=15-20). **(F)** Western blot lanes# **1:** Fv, reproductive tract (RT) of 5 virgin females (negative control) **2:**, M, a pair of male accessory gland (positive control), **3**,**4**,**5:** RT of females mated to CS males, flash frozen at 2hrs (n=5), 1d (n=15) and 4d (n=15) ASM respectively, **6**,**7**,**8**: RT of females mated to *SP*-null males and then subsequently mated to spermless males at 1d ASFM, flash frozen 2hrs (n=5), 1d (n=15) and 4d (n=15) ASSM, respectively. Actin served as loading control. **(G)** Sperm in the seminal receptacle (SR) of a typical females mated to a control male (*SP*+; *ProtB-eGFP*) at 4d ASM. **(H)** Sperm in the SR of a typical female mated to *SP-null;ProtB-eGFP* male at 4d ASM. **(I)** Sperm in the SR of a typical female, mated to *SP-null; ProtB-eGFP* and subsequently remated to a spermless male at 1d ASFM, and frozen at 4d ASSM. In G-I sperm heads are green due to eGFP. Bar=50µm. **(J)** Graphical representation of sperm counts in SRs of females singly-mated to control (SP+, red, TM3 siblings of *SP-null; ProtB-eGFP*), *SP*-null (green) or doubly-mated to *SP*-null and spermless male (*SP-*null, SOT, blue) represented in G, H, I panels (p**=<0.01; p*=<0.05; ns=non significant; n=15-20).

Reducing the likelihood of mated females to remate is another crucial postmating response regulated by SP [35,37]. Females that do not receive SP generally fail to exhibit this reluctance, and remate readily. We tested whether SP from a second male could delay the receptivity of females that had previously mated to *SP*-null males. Females singly-mated to *SP*-null males or spermless males show a significantly higher tendency to remate at 1d ASM (Fig 3D; p***=<0.001) or 4d ASM (Fig 3E; p***=<0.001) relative to females mated to wt (CS) males (Fig 3D and 3E). In contrast, females mated to *SP*-null males and then remated to spermless males at 1d ASFM (Fig 3D; p=0.43) showed receptivity similar to mates of control males at 1d after the start of second mating (ASSM). The effect, however, did not persist as long as after a mating to a wt male. At 4d ASSM (Fig 3E; p***=<0.001) doubly-mated females exhibited higher receptivity relative to females mated to wt males but lower than those mated to spermless males. This could be either because less SP from the second (spermless) mating is able to bind to stored sperm from the previous mating and thus SP levels have been more depleted by 4 days ASSM than after a control mating where the sperm-SP enter the female together. Alternatively, the active portion of SP received from a rival male, bound to first male’s sperm might be released from the sperm at a higher rate. We performed western blotting to determine how long SP received from the second (spermless) male persists in the reproductive tract of females previously mated to *SP*-null males. Females singly-mated to CS males and those doubly-mated to *SP*-null males and spermless males at 1d ASFM, were flash frozen at 2hr, 1d or 4d ASM/ASSM, respectively. SP signals were detected in females mated to CS males at 2hr, 1d or 4d ASM (Fig 3F. lanes 3, 4, 5). SP was detected in females mated to *SP*-null males and then remated to spermless males at 2hr and 1d ASSM (Fig 3F. lanes 6, 7) but not (or very weakly) at 4d ASSM (Fig 3F. lane 8). Taken together, our results show that SP from a second male can rescue the receptivity defects that resulted from the first male’s of lack of SP but that sufficient SP for such an effect is not retained for as long as in a control situation (e.g. a mating with a wt male).

SP is also needed for release of sperm from storage within the mated female [6]. Thus, females mated to *SP*-null males retain significantly more sperm in their seminal receptacle at 4d ASM. To test whether SP acquired from a spermless male in a second mating could also rescue this defect, we counted sperm in storage after a single mating with *SP-null*; *ProtB-eGFP* males and after mates of *SP-null*; *ProtB-eGFP* males had remated with spermless males. As expected, females mated to control (*SP*^+^; *ProtB-eGFP*) males had fewer sperm in their seminal receptacle (average of 192; Fig 3G and 3J) relative to mates of *SP-null*; *ProtB-eGFP* males, which had significantly higher sperm counts, indicating poor release of stored sperm (Fig 3H and 3J; p**=<0.01; average of 304 at 4d ASM). However, mates of *SP-null*; *ProtB-eGFP* males that had remated with spermless males retained sperm in numbers similar to those observed in females mated to control males (average of 212; Fig 3I and 3J; p=ns). We also counted sperm stored in seminal receptacle of females mated to *SP-null*; *ProtB-eGFP* males at 5d ASM (Fig 3J) to make sure that the evident decline in sperm counts or release of stored sperm in doubly mated females (*SP-null; ProtB-eGFP* mates remated to spermless males that were frozen at 5d ASFM or 4d ASSM) was not dependent on days after mating, rather on receipt of SP from spermless males. Thus, SP from a second male can rescue the sperm release defects of prior matings to males that lacked SP.

### 3. SP from a second male can bind to stored sperm from a previous male, while still binding strongly to his own sperm

In the experiments described above SP was provided by a spermless second male, but in nature females are much more likely to encounter a male who has his own sperm, capable of binding his SP. To test whether SP from a male with sperm can still bind to sperm from another male, we modified our experimental protocol such that females were mated to *SP*-null males as described earlier, but rather than spermless males, we now used *ProtB-dsRed* males [38] as the second male (Fig 4I. Cartoon). These second males have a full suite of SFPs, sperm and their sperm-heads are labeled with ProtB-dsRed. This allowed us to distinguish between sperm received from *SP*-null males (blue heads) and those received from *ProtB-dsRed* males (red heads). Females were frozen at 2hrs ASSM and sperm dissected from their seminal receptacles were probed for SP. We observed anti-SP staining along the entire sperm (head and tail) from *ProtB-dsRed* males (Fig 4B). Sperm received from the *SP*-null males (blue heads) were also stained with anti-SP along their length (head and tail; Fig 4B). Therefore, a control (wt) male with a complete suite of SFPs and sperm of his own can also provide SP to bind to SP-deficient sperm from another male.

**Figure 4.**
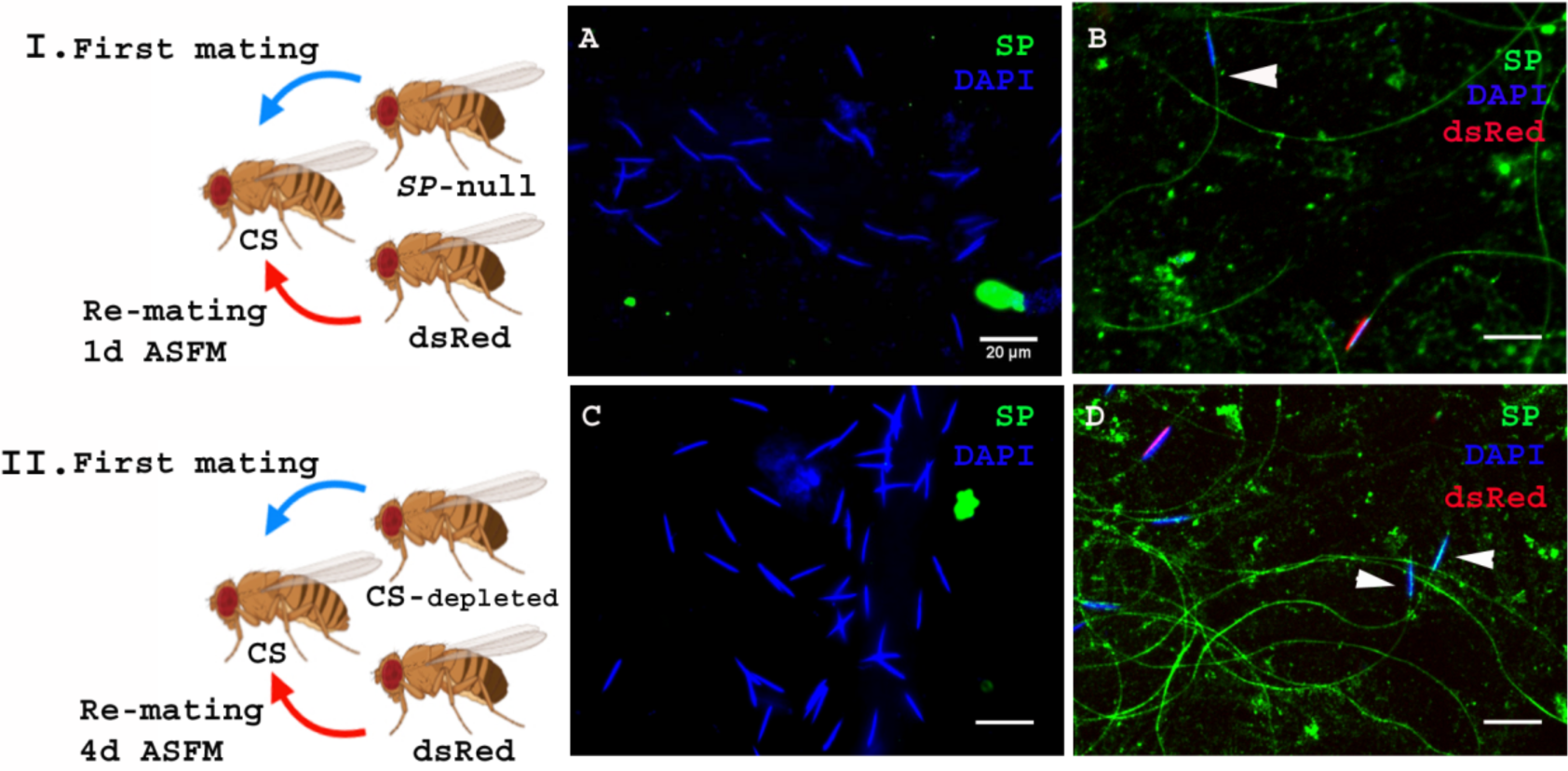
SP from a male who also provides sperm can bind to SP-deficient sperm as well as to the donor’s sperm. **Cartoon (I):** Pictorial representation of the experimental cross (fly images from Biorender). Females mated to *SP*-null males were remated to control (*ProtB-dsRed*) males at 1d ASFM. **(A)** Sperm from females singly mated to *SP*-null males, 2hr ASM (blue sperm-head). **(B)** Sperm from females mated to *SP*-null males (blue sperm-head) remated to *ProtB-dsRed* (red+ blue sperm-head) males at 1d ASFM. SP visualized with Alexa fluor 488, staining the sperm (head+ tail; green). Flies were frozen 2h ASSM. White arrows indicate sperm heads (n=10; Bar = 20µm). **Cartoon (II):** Pictorial representation of the substitute cross (fly images from Biorender). Females mated to SFP depleted control (CS) males were remated to control (*Prot B-dsRed*) males at 4d ASFM. **(C)** Sperm from females singly-mated to SFP depleted CS males at 4d ASM (blue sperm-head). **(D)** Sperm from females mated to SFP depleted CS males (blue sperm-head), remated to *ProtB-dsRed* (red+ blue sperm-head) males at 4d ASFM. SP visualized with Alexa fluor 488, staining the sperm (head+ tail; green). Flies were frozen 2h ASSM. White arrows indicate sperm heads (n=10; Bar = 20µm).

The likelihood of finding an *SP*-null male in nature is very low. However, multiple-mating has been shown to deplete SFP reserves [39], so it is possible that inter-ejaculate interaction (or copulation complementation) could occur if the first male had depleted his SFP reserves. To test whether this could occur, we performed a crossing scheme in which we substituted multiply-mated control (CS) males with exhausted seminal reserves [39] for the *SP*-null males used in Fig. 4A. We carried out western blotting to determine the levels of SP in accessory glands (AG) of such multiply mated (CS) males and the amount of SP in their mates at 2hr ASM. We observed relatively weak SP signals in the AG of multiply-mated males (Fig S2. A, lane 4) and a very faint SP signal in females mated to these males (Fig S2. A, lane 5) compared to relatively strong SP signal in virgin (unmated) males and the females mated to these males (Fig S2. A, lane 2, 3 respectively). Our immunofluorescence data showed no (or extremely weak) SP-sperm binding in sperm dissected from the seminal receptacle of females mated to SFP-depleted males (Fig S2. C). Females mated to SFP-depleted CS males were then subsequently remated at 4d ASFM (long enough to have lost any SP signal from their first multiply-mated, mates) to *ProtB-dsRed* males. Sperm were dissected from the seminal receptacles of these females at 2hrs ASSM, and probed for SP (Fig 4II. Cartoon). We did not observe any detectable SP signal on sperm stored in females singly-mated to SFP-depleted CS males at 4d ASM (Fig 4C). However, we observed anti-SP staining along the entire sperm (head and tail) received by the doubly-mated female from the SFP-depleted CS male (blue heads; Fig 4D) and *ProtB-dsRed* males (red+ blue heads; Fig 4D).

Thus, in a normal mating the amount of SP that a male transfers is sufficient to bind not only his own sperm but also to remaining sperm from a rival. Moreover, SP from an unmated control male can bind to previously stored sperm of a male that had his SFP reserves depleted prior to mating with the female.

### 4. Sex peptide binding to sperm of a prior male does not require receipt of LTR-SFPs from the second male

SP binding to sperm requires the action of a network of other SFPs- “LTR-SFPs” [10,21]. Most of the known LTR-SFPs bind to sperm transiently (CG1656, CG1652, CG9997 and antares) [20], while others do not bind to sperm (CG17575 or seminase) [40]; the latter facilitate the localization of other LTR-SFPs, and SP, to the seminal receptacle. However, no LTR-SFPs are detectable on sperm or in female RT at 1d ASM (Fig 5 & Fig S3). We wondered whether LTR-SFPs were required from the second male in order to bind his SP to the first male’s sperm.

**Figure 5.**
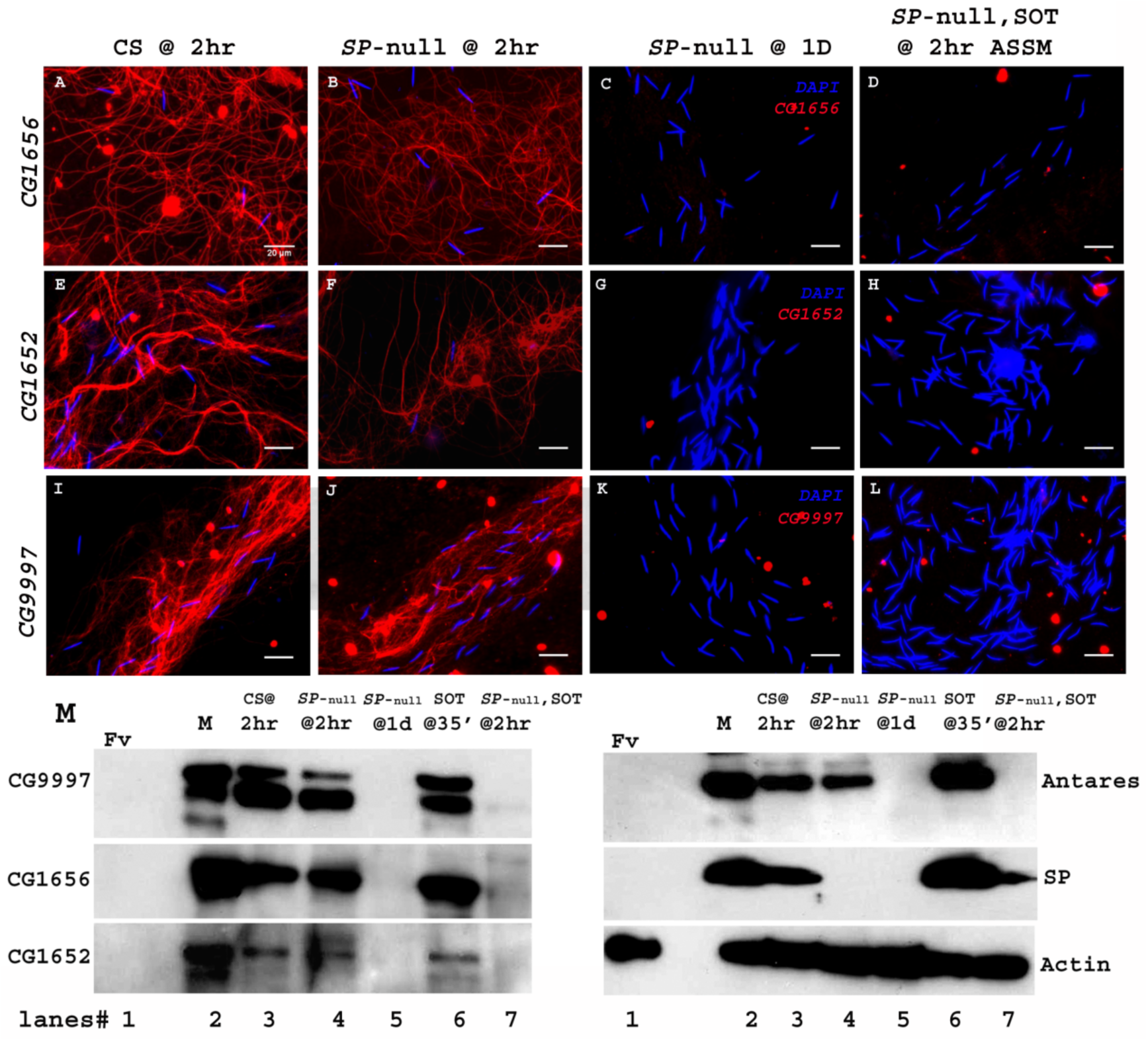
Sperm do not bind detectable LTR-SFPs from a second male. Females mated to wild type (CS) males at 2hr ASM show LTR-SFPs bound to sperm, CG1656 **(A)**, CG1652 **(E)**, CG9997 **(I)**. Females mated to *SP-*null males show the same **(B**,**F**,**J)** but by 1d postmating LTR-SFPs’ signal were no longer detected on sperm **(C**,**G**,**K)** confirming previous reports (please see [20]). Females mated to *SP*-null males and then remated to spermless males also do not show detectable signal for sperm-LTR-SFP binding for CG1656 **(D)**, CG1652 **(H)** and CG9997 **(L)**, 2hrs ASSM, although they have SP bound (Fig 1). Sperm stained for the indicated LTR-SFP detected with Alexa fluor 594 (red) and sperm-head stained with DAPI (blue). Bar=20µm **(M)** Western blot probed for indicated LTR-SFPs. Lanes/samples are, **1:** Fv, reproductive tract (RT) of 3 virgin females (negative control), **2:** M, 1 pair of male accessory glands (positive control), **3:** CS@2hr, sperm dissected from SR of 20 females mated to wild type (CS) males at 2hr ASM, **4:** *SP*-null @ 2hr, sperm dissected from SR of 20 females mated to *SP*-null males at 2hr ASM, **5:** *SP*-null @1d, sperm dissected from SR of 20 females mated to *SP*-null males at 1d ASM, **6:** SOT@35’, reproductive tract of 3 females mated to spermless males at 35’ASM (positive control), **7:** *SP*-null, SOT @ 2hr, sperm dissected from SR of 20 females mated to *SP*-null males and then remated to spermless males at 1d ASFM, and frozen at 2hr ASSM. Lanes were probed for LTR-SFPs CG9997, CG1656, antares and CG1652 and SP as described in the text. Actin served as loading control.

We carried out experiments similar to those previously described, in which females were first mated to *SP-*null males and then remated to spermless males at 1d ASFM. We froze females at 2hr ASSM and immunostained stored sperm dissected from their seminal receptacles, for the presence of LTR-SFPs that had been received from second (spermless) males.

Females mated to CS males and frozen at 2hr ASM served as positive controls for the sperm-binding of LTR-SFPs, CG1656 (Fig 5A), CG1652 (Fig 5E) and CG9997 (Fig 5I). Females singly mated to *SP*-null males, frozen at 2hr ASM, exhibited normal sperm-binding of LTR-SFPs, CG1656 (Fig 5B), CG1652 (Fig 5F) and CG9997 (Fig 5J), confirming that loss of SP affects neither the transfer nor the sperm-binding of other LTR-SFPs [20]. By 1d ASM, stored sperm from females singly-mated to *SP-*null males showed no signal for the LTR-SFPs, CG1656 (Fig 5C), CG1652 (Fig 5G) and CG9997 (Fig 5K), as expected given the transient sperm-binding of these proteins [20]. Thus, by the time these females remated with spermless males (1d ASFM), all known LTR-SFPs received from the first (*SP*-null) male were undetectable on sperm.

Interestingly, although females that mated to *SP*-null males and then to spermless males showed SP signal on their sperm (as in Fig 1) at 2hrs ASSM, we detected no signal of LTR-SFPs, CG1656 (Fig 5D), CG1652 (Fig 5H) and CG9997 (Fig 5L) on those sperm at 2hrs ASSM. This could be because LTR-SFPs from the second male could not enter the sperm storage organs in the absence of sperm or, alternatively, that their binding sites on sperm had been modified prior to the second mating (perhaps by the action of LTR-SFPs received from the first mating) to make them incapable of binding.

We verified these observations by western blots. Consistent with immunofluorescence data as in Fig 5A-L, LTR-SFP signals for CG1656, CG9997, CG1656 and Antares were detected in sperm dissected from females mated to CS and *SP-*null males at 2hr ASM (Fig 5M. lanes 3, 4). No LTR*-*SFP signals were detected in sperm dissected from females mated to *SP*-null males at 1d ASM (Fig 5M. lane 5) or in sperm dissected from females mated to *SP*-null males, remated to spermless males at 1d ASFM, and frozen 2hrs ASSM (Fig. 5M. lane 7). However, as expected, SP signals were detected in sperm dissected from females that mated to *SP*-null males, remated to spermless males at 1d ASFM and frozen 2hrs ASSM (Fig 5M. blot probed for SP, lane 7).

Thus sperm no longer detectably bind new LTR-SFPs after they have bound LTR-SFPs from their own (*SP-*null) male. That LTR-SFPs are needed for SP-sperm binding, and that SP from spermless male binds the first male’s sperm, further suggests that the first male’s sperm (or the female RT) had already been primed with its own LTR-SFPs during storage in the female tract.

Unlike the four LTR-SFPs assessed above, the two other LTR-SFPs, CG17575 and seminase, do not bind to sperm, yet are crucial for SFP-sperm binding. In the absence of CG17575 or seminase, SP fails to bind to sperm [10,40]. To determine if these proteins were required for a second male’s SP binding to a first male’s sperm, we first crossed females to *SP-*null males and then to *CG17575*-null or *seminase*-null males at 1d ASFM (Fig 6. Cartoon). In this situation, CG17575 and seminase had entered the female with the first male’s sperm, but by the time of the second mating, were undetectable in the female (Fig S3). We examined whether in this situation SP transferred by *CG17575*-null (or *seminase*-null) males would still bind to the *SP-*null sperm stored in the female. We made use of ProtB-eGFP labelled *SP*-null males to differentiate between sperm received from first (cyan (DAPI+ eGFP) sperm heads) and second (blue (DAPI) sperm heads) males. Immunostaining and western blots for detection of SP on sperm dissected from females mated to *SP-null; ProtB-eGFP* males and then remated to *seminase*-null (Fig 6. A and C, lane 4) or *CG17575*-null (Fig 6. B and C, lane 5) males showed that SP received from the second male bound to sperm (head and tail) received from *SP-null; ProtB-eGFP* males. Sperm dissected from females singly-mated to *SP-null; ProtB-eGFP* males gave no SP signal, as expected (Fig 6. D, lane 3 and E) and sperm dissected from females singly-mated to *seminase*-null (Fig 6. D, lane 4 and F) or *CG17575*-null (Fig 6. D, lane 5 and G) males also showed no SP-sperm binding, as expected, due to lack of the LTR-SFP. Therefore, sperm no longer require even CG17575 or seminase from the second male’s ejaculate, after they have received the LTR-SFPs from their own (*SP-*null) male.

**Figure 6.**
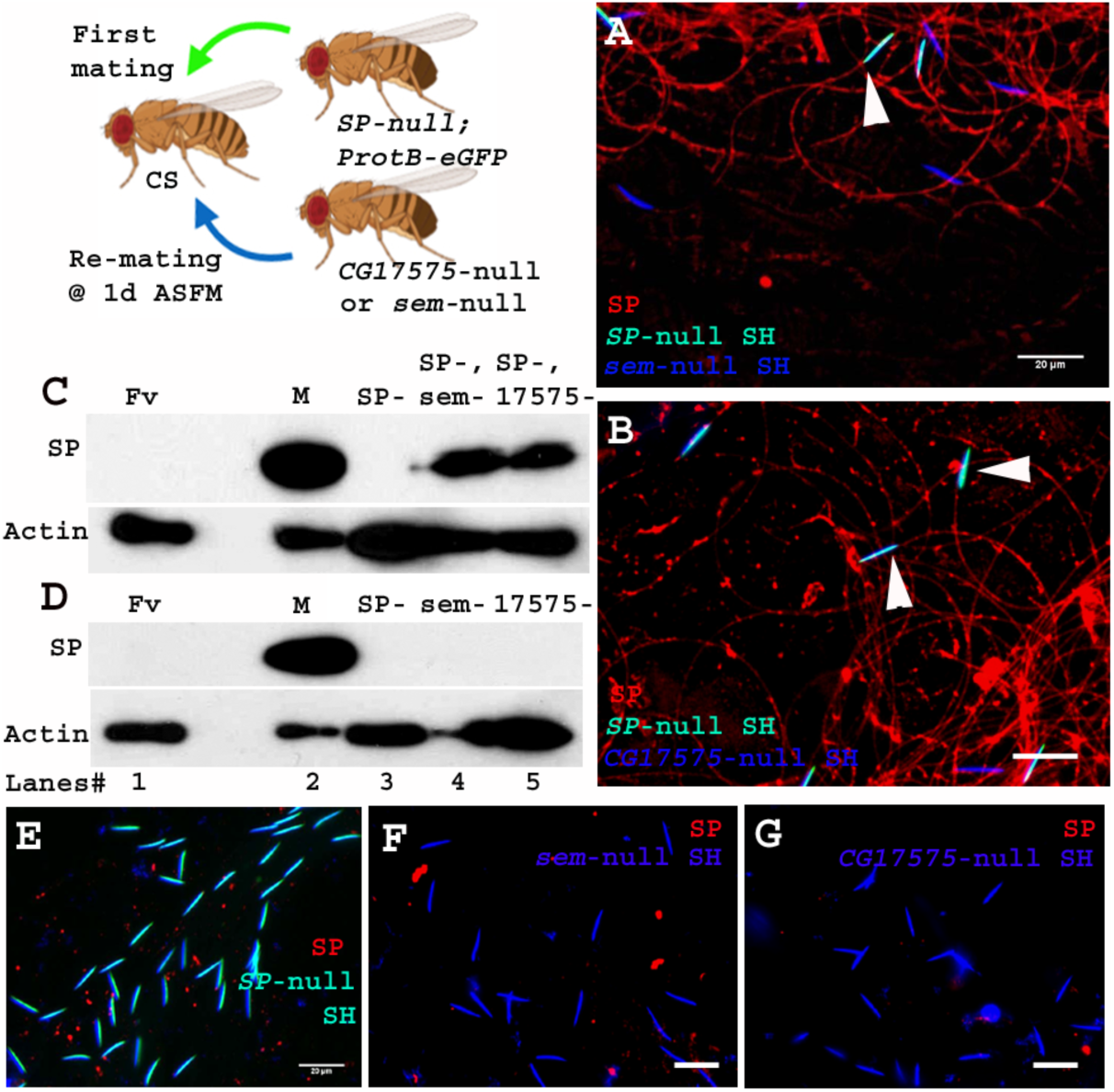
Sperm received from SP-null males do not require CG17575 or seminase from a second male to bind SP from that male. **Cartoon:** Pictorial representation of the experimental cross (fly images from Biorender). Females mated first with *SP-null; ProtB-eGFP* male [cyan sperm-head; DAPI(blue)+eGFP(green)] and then a day later with *CG17575-*null or *seminase*-null male (blue sperm-head; DAPI stained) and frozen, 2hrs ASSM. SP was visualized with Alexa fluor 594, staining the sperm (head+ tail) red. **(A)** Sperm from females mated to *SP-null; ProtB-eGFP* males and then remated to *seminase-* null males, 1d ASFM. **(B)** Sperm from females mated to *SP-null; ProtB-eGFP* males and then remated to *CG17575-*null males, 1d ASFM. **(C)** Western blot probed for SP. Lanes/samples are, **1:** Fv, reproductive tract (RT) of 3 virgin females (negative control), **2:** M, 1 pair of male accessory glands (positive control), **3:** SP-, sperm dissected from 20 females mated to *SP-null; ProtB-eGFP* males at 2hr ASM, **4:** SP-, sem-, sperm dissected from 20 females mated to *SP-null; ProtB-eGFP* males and subsequently to *seminase*-null males at 1d ASFM, frozen at 2hrs ASSM, **5:** SP-, 17575-, sperm dissected from 20 females mated to *SP-null; ProtB-eGFP* males and subsequently to *CG17575*-null males at 1d ASFM, frozen at 2hrs ASSM. **(D)** Western blot probed for SP. Lanes/samples are, **1:** Fv, reproductive tract (RT) of 3 virgin females (negative control), **2:** M, 1 pair of male accessory glands (positive control), **3:** SP-, sperm dissected from 20 females mated to *SP-null; ProtB-eGFP* males at 2hr ASM, **4:** sem-, sperm dissected from 20 females mated to *seminase*-null males at 2hr ASM **5:** 17575-, sperm dissected from 20 females mated to *CG17575-*null males at 2hr ASM. Actin served as loading control. **(E)** Sperm isolated from females singly mated to *SP-null; ProtB-eGFP* males, 2hr ASM. **(F)** Sperm isolated from females singly mated to *seminase*-null male, 2hr ASM. **(G)** Sperm isolated from females singly mated to *CG17575*-null male, 2hr ASM. White arrows indicate sperm heads (represented as SH, n=10; Bar=20µm).

## Discussion

Ejaculate molecules, particularly the seminal fluid proteins (SFPs) that are received by females during mating, play crucial roles in successful fertilization. In *Drosophila* they induce striking changes in the physiology and behavior of females, instigating a wide array of post mating responses [6,14,28,29,40–42]. Some of these responses persist long-term, due to binding of a male’s SP to his sperm and gradual release of the SP’s active C-terminal region [19]. This important process is mediated by a cascade of “LTR-SFPs” that are needed to bind SP to sperm [10,20,21,40]. While all of the above can be seen as facilitating reproductive success of the mating pair (particularly from the male’s perspective), SFPs also play roles in conflicts between males in species where females are polyandrous. Den Boer et al [43] investigated sperm survival in monandrous and polyandrous ants and bees. They observed that while seminal fluid enhanced the survival of “self” sperm, it preferentially killed the sperm of rival males. In other words, while SFPs worked in a cooperative interdependent way with “self” sperm, they harmed rival sperm when in a situation of conflict (and cryptic female choice). Previous studies have shown that males respond to threat of rivals by altering the allocation of both sperm as well non-sperm components of their ejaculate (e.g for *Drosophila*: [11,25,26]).

Studies of SFP functions have tended to investigate how a male’s SFPs can promote the interests of his own (self) sperm. However, some data suggest that one male’s SFPs (ovulin, ACP36DE) can indirectly benefit a subsequent male within a polyandrous female [27–30,44]. Here, we tested for direct effects of one male’s SFPs on another male’s sperm and/or fertility. Specifically, we show that SP from a second male can bind to and act with sperm received from a previous mating. Sperm stored in females mated to *SP*-null males show no SP-sperm binding (as expected), but if these mated females subsequently remate to a spermless male, his SP can bind to stored sperm from the prior male. This binding of SP to the *SP*-null sperm restores his fertility and proper sperm release dynamics. Even if a second male transfers sperm, he transfers sufficient SP to bind to his own and rival sperm. Finally, our data suggest that the LTR-SFPs (that usually assist in binding of SP to sperm) are not required from the second male for the association of his SP with sperm received from the first male (who had already provided LTR-SFPs). The first male’s sperm appear to be sufficiently “primed” by prior receipt of their own LTR-SFPs to be able to bind SP from a second male.

### SP from a second male can associate with a prior male’s sperm that were stored within the female

Xue and Noll [31] reported that sperm transferred to females by *Prd* mutant males (that lack the entire suite of SFPs) were capable of fertilizing a few eggs to yield progeny, but only after the females were subsequently remated to spermless males. They coined the term “copulation complementation” to describe this phenomenon, and proposed that SFPs from the second male might interact with the first male’s sperm to yield this result. Consistent with the idea of copulation complementation, several reports suggested that first-males that lacked particular SFPs [6,8,45–47] can have reproductive advantage in terms of their higher paternity share in competitive situations; they were better competitors, compared to controls, in defensive sperm competition assays [6,29,48]. A simple explanation for these results, based on the findings that we report here, is that the deficiency in the first male led to impaired release/use, and thus retention, of his sperm, and this was complemented by receipt of the second male’s SFPs, as we have now shown here, for SP.

SP is the only SFP thus far known to persist within the *Drosophila* female (for 10-14 days post-mating), eliciting long-term post mating responses through gradual release of its C-terminal portion [19]. The long-term persistence of SP on sperm made it an excellent candidate to examine for interaction with rival sperm. Here, we report that SP subsequently received from a spermless male binds to a first male’s sperm (*SP*-null). This association is apparent even if the second mating occurs at 1d or as long as 4d ASFM, indicating that binding of SP to the first male’s sperm occurs irrespective of how long sperm have been in the storage organs. It remains unclear how SP received from spermless (second) male enters the sperm storage organs, where sperm from the first mating had been stored. However, Manier et al [38] reported that 60-90 min after the start of a second mating, 26% of the resident sperm (received from the previous mating) are moved from storage back into the bursa where they mix with the second male’s ejaculate before moving back into storage. Therefore, it is possible that SP received from the spermless male binds to the first male’s sperm that relocated to the bursa, and the newly SP-bound sperm are then transferred back into storage in the seminal receptacle.

### The binding of SP received from one male to sperm of another can restore defects that resulted from lack of SP from the first male

In the absence of sperm, or if SP is not bound to sperm, females do not maintain post-mating responses and fail to efficiently release sperm from storage resulting in fewer sperm available for fertilization and fewer progeny [6,10,17,35]. We observed that these defects were restored when SP was received by females in a remating with spermless males. Thus, the second male’s SP bound to the first male’s sperm is functional. The rescue of the phenotype, however, was not as long lasting as in a normal single mating with SP transfer, wearing off by 4d postmating rather than the normal ∼10d. This could be because only fewer sperm relocated from storage to the bursa [38], so they may not carry sufficient SP back into storage to associate with *SP*-null sperm. Consistent with this, the levels of SP that we see stained in these situations are lower than those in a wild type mating.

### An unmated male transfers sufficient SP to bind to his own as well rival sperm

We did not know whether the amount of SP that is transferred during mating is more than the available binding sites on sperm. Here, we observed that an unmated control male does transfer enough SP to bind his own as well as pre-stored sperm (*SP*-null) in a previously mated female. Consistent with our findings, several reports suggest that in response to potential threats of sperm competition and conflicts, males allocate the levels of SFPs and transfer more SP, yet less ovulin, to previously mated females [11,26].

Rubinstein et al [28] demonstrated that ovulin induces ovulation, acting through octopamine (OA) neuronal signalling and increases the number of synapses that the female’s Tdc2 neurons make on the musculature of the oviduct. This latter effect persisting could benefit rivals too, so males may thus be able to mitigate the levels of ovulin in their ejaculate. But the question remains that if SP from one male’s ejaculate can bind to and assist another’s sperm, why do males not lower the amount of SP transferred while mating? A potential explanation is that a male would still benefit by transferring enough SP to ensure that his own sperm remains saturated with SP, even at a cost of part of his SP binding to another male’s sperm.

SP binds to sperm through its N-terminal region, and this region remains bound to sperm long-term [19]. The bound N-terminal region of SP on sperm stored in a mated female does not allow any further binding of SP coming from rival male’s ejaculate. Therefore under what circumstances might SP-mediated copulation complementation occur in nature? In polygynous males, SFPs are depleted faster than sperm [39]. This could result in a situation in which a female who mated with a male with low levels of SFPs might not receive enough SP to saturate his sperm. In these circumstances, SP received from another male would help compensate for the lower amount of SP from the depleted first male’s ejaculate. SFP depletion would, of course, not only affect the levels of SP, but also all the other crucial LTR-SFPs. However, while other LTR-SFPs enable SP to bind sperm, it is the quantity of bound SP that correlates with the duration of post-mating responses. In line with this hypothesis, we subjected a control male to recurrent matings (providing six virgins) over the span of two days, with an intent to exhaust their SFPs. We observed that sperm stored by subsequent (7^th^) females mated to these multiply-mated males had undetectable SP signals. However, when these females were remated to unmated control males, strong SP signals were detected on both the SFP-depleted sperm received from the previous mating and the newly received rival sperm.

Therefore, our results support the idea that in nature males who have multiply-mated might get some help from the SFPs of subsequent, less depleted, males. Interestingly, this inter-ejaculate interaction might confer an added advantage to the second male. More of the second male’s SP will be retained in the female reproductive tract, for even longer, if it binds to previously-stored sperm in addition to his own sperm. This could allow the post-mating responses in polyandrous females to be maintained for longer than in singly-mated females.

### Association of a second male’s SP to sperm received from a prior male does not require the receipt of LTR-SFPs from the second male

Binding of SP to sperm is facilitated by a network of LTR-SFPs [10]. Two LTR-SFPs, CG17575 and seminase, do not themselves bind to sperm, whereas other LTR-SFPs bind sperm transiently (CG1652, CG1656, CG9997, antares). CG17575 and seminase localize the other LTR-SFPs, and SP, to sperm storage organs [10,20,40,42,49]. We found that SP from a second male (spermless or control) can associate with sperm from the first male (*SP*-null) even if it enters the female in absence of its own LTR-SFPs. This suggested that *SP*-null sperm (or the mated female RT) had already received modifications (“priming”) from its own LTR-SFPs that were required for SP binding. This further suggests that once primed, a sperm can bind SP from a rival’s ejaculate without the need for additional LTR-SFPs, and can restore its own post-mating dynamics.

Thus, we find that a critical SFP from one male can associate and offer direct benefits to sperm from another male, restoring the SP function to the previously stored sperm. Our work shows that SP is a crucial candidate for copulation complementation in *Drosophila*, and that sperm in storage (or the female RT) are primed for SP binding by the first male’s LTR-SFPs. Thus, despite potential competition between males, there could be subtle cooperation between males as well. In addition, the allocation of resources by, and effects on, rival males that mate to polyandrous females, should be viewed in light of not only sexual conflicts, but also both direct and indirect effects of SFPs.

## Materials and Methods

### 1. Fly strains

Spermless males, [*sons of tudor*, (*sot*) that lack sperm but produce and transfer a complete suite of SFPs] were the progeny of *bw sp tud*^*1*^ females [50] mated to control, *Canton S* (CS) males. *Sex peptide* null mutant males (*Δ325/Δ130*; which have sperm and the entire suite of SFPs except for SP) [35] were generated by crossing the SP knockout line (*Δ325/TM3, Sb ry*) to a line carrying a deficiency for the *SP* gene (*Δ130/TM3, Sb ry*). Control males were the TM3 siblings of *SP*-null mutants. Matings were conducted with wild type *D. melanogaster* females (CS). To determine sperm numbers, we generated a line carrying the *SP*-null mutation and Protamine B-eGFP tagged sperm (*ProtB-eGFP/Y; Δ325/Δ130*) by series of crosses between the *SP* knockout line (*Δ325/TM3, Sb ry*) and *ProtB-eGFP (X); TM3/TM6* [38]. The *TM3* siblings of these males, (*SP*^+^; *ProtB-eGFP)* served as controls. Sperm-heads of these control males were tagged with ProtB-eGFP, but the males had normal levels of SP (Fig S4). All flies were reared under a 12:12h light-dark cycle at 22±1°C on standard yeast-glucose medium. Mating experiments were carried out by single-pair mating 3-5 day old virgin CS females to 3-5 day old unmated males of genotypes indicated in the text and remating the same female 1 day or 4 days after the start of first mating (ASFM) to age matched unmated males of the genotypes indicated in the text.

### 2. Crossing scheme to study first male’s sperm and rival’s SP binding

Xue and Noll [31] reported copulation complementation in females mated to *Prd* males (which produce sperm but lack SFPs) remated to spermless males (*sot*, which produce SFPs). We followed a similar scheme but to focus on SP specifically, we used *SP*-null males as the first male. As described in Results, we then remated these females to spermless males, which make SFPs but not sperm. We attempted to do the reciprocal experiment, where females were mated to spermless males and then remated to *SP*-null males, but consistent with what was reported by Xue and Noll [31], we could not detect copulation complementation in this direction for technical reasons: SP from the spermless male did not persist long enough in the mated female to interact with the second male’s sperm (see Results). We carried out rematings at three time points, 3-6 hrs, 1d, and 4d AFSM. We assessed results at 2hr after the start of the second mating (ASSM).

### 3. Fertility

The reproductive performances of singly-mated or doubly-mated females were assayed by analyzing fertility (numbers of progeny eclosed over ten days) [32]. Briefly, the fertility assays were carried out with (A). “Single matings”: Females were singly mated to (i) spermless males, (ii) *SP*-null males, or their (iii) *TM3* siblings (genetically-matched control males) in three individual sub-batches, and (B). “Rematings”: Females were mated to *SP*-null males or their *TM3* siblings (*SP*^+^) and were then subsequently remated to spermless males at 1d and 4d ASFM. Matings that lasted 15 mins or more were considered successful. At the end of a mating, males were removed from the vials and females were allowed to lay eggs for 10 days after the start of mating (ASM) in the first batch and after the start of second mating (ASSM) in the second batch. Females were transferred to fresh food vials every three days. Flies emerging from each vial were counted. Fertility is represented as total progeny number produced by each female over a period of 10 days. The differences in fertility were analyzed through One way Analysis of Variance (ANOVA) followed by Tukey’s multiple comparison tests for single-matings and Mann Whitney U-tests for rematings. All assays were repeated more than two times and comprised of two technical replicates, with each group consisting of a minimum sample size of 15-20.

### 4. Receptivity

To determine the propensity of females to remate, receptivity assays [17] were set for females singly mated to *SP*-null, spermless or CS males and females mated to *SP*-null males and then subsequently remated to spermless males at 1d ASFM. For the assay, females from singly-mated and doubly-mated groups were then provided with (CS) males at 1d and 4d ASM or ASSM, respectively. We determined the number of females that mated within 1hr from when the CS male was introduced within the vial. Assays were repeated more than two times, with each group consisting of a minimum sample size of 15-20. The data were analyzed by Fisher exact tests and Chi-squared group analyses.

### 5. Sperm utilization/ release from sperm storage organs in females

To study the effect of first male’s sperm and rival male’s SP binding on sperm utilization and release, we generated *SP*-null males whose sperm-heads are labelled with ProtB-eGFP [38]. Females were mated to *SP-null*; *ProtB-eGFP* or *SP*^+^; *ProtB-eGFP* (control) males. Some of the mated females were frozen at 4d ASM (or 5d ASM) for sperm counts. The remaining mates of *SP-null*; *ProtB-eGFP* males were remated to spermless males at 1d ASFM. These flies were flash-frozen at 4d ASSM. Subsequently, seminal receptacles of females singly-mated to *SP-null; ProtB-eGFP* and *SP*^+^; *ProtB-eGFP*, or doubly-mated to *SP-null; ProtB-eGFP* and spermless males, were dissected and eGFP sperm were counted (at a total magnification of 200X, with FITC filter on an Echo-Revolve microscope). Mature sperm in the seminal receptacles of mated females were counted twice and groups were blinded to ensure reproducibility and avoid bias. The percent accuracy was 90-94%. Assays were repeated more than two times, with two technical replicates. Every group contained a minimum sample size of 15-25. Differences in the sperm counts between groups were analyzed statistically through One way ANOVA followed by Tukey’s multiple comparison tests.

### 6. Brood matings

Control (CS) males were subjected to brood matings [51,52] to deplete SFPs, as their levels are known to become exhausted at a higher rate than sperm numbers [39]. Briefly, three day old control males were mated to CS females in two broods (each consisting of three virgin females) over two days. The first mating of both broods was observed. On the third day, previously mated females were removed and the male was provided with an additional virgin female (7^th^ mate), matings were observed and depleted CS males were removed. Half of the 7^th^ mated females were frozen at 4d ASM, while the others were subsequently remated to control (*ProtB-dsRed*) males at 4d ASFM, and then frozen at 2hr ASSM. Sperm stored in the seminal receptacle of the frozen flies were dissected and immunostained for SP.

### 7. Immunofluorescence

Immunofluorescence was performed to detect SP-sperm binding [10,19,20]. Sperm dissected from seminal receptacles of experimental or control females were attached to poly-L-Lysine (Sigma) coated slides. Sample processing was carried out according to the protocol of Ravi Ram and Wolfner [10] with minor modifications. Samples were blocked with 5% bovine serum albumin, BSA in 0.1% PBX for 30min. Subsequently, samples were incubated overnight in rabbit anti-SP(1:200), CG1656(1:100), CG1652(1:50), CG9997(1:50) [20], in 0.1%BSA at 4°C overnight. Samples were then washed in PBS and incubated at room temperature for 2h in mouse anti-rabbit IgG coupled to alexa fluor 488 (green) or 594 (red; Invitrogen) at a concentration of 1:300 in 1xPBS at room temperature in the dark. Samples were then washed in PBS, incubated in 0.01% DAPI for 3 min at room temperature in the dark, rewashed and mounted using antifade (0.2% N-propyl gallate in 75% glycerol; Sigma). The fluorescence was visualized under an Echo-Revolve fluorescence microscope at a magnification of 200X. A minimum of three independent immunostaining batches, with a minimum sample size of 10, were analyzed for each group.

### 8. Sample preparation and Western blotting

To further examine transfer, persistence or binding of SP to sperm stored in singly-mated or doubly-mated females, the lower reproductive tract (RT) or sperm stored (SS) in seminal receptacles of mated female were dissected. The dissected tissues (lower RT, n=5-10 or sperm, n =20-30) were suspended in 5µl of homogenization buffer (5% 1M Tris; pH 6.8, 2% 0.5M EDTA) and processed further according to the protocol of Ravi Ram and Wolfner [10]. Proteins from stored sperm or lower female reproductive tract were then resolved on 12% polyacrylamide SDS gel and processed further for western blotting. Affinity purified rabbit antibodies against SP(1:2000), CG1656(1:1000), CG1652(1:500), antares(1:500), CG9997(1:1000), CG17575(1:1000), seminase(1:1000) [10,20,40] and mouse antibody against actin (as a loading control; Millipore Corp., cat no. #MAB1501MI at 1:3000) were used as primary antibodies. HRP conjugated secondary anti-rabbit and anti-mouse antibodies (Jackson Research) were used for detection of SFPs at a concentration of 1:2000.

## Acknowledgements

We thank Dr. Ravi Ram Kristipati, S. Allen, N. Brown, S. Delbare and D. Chen for helpful suggestions and comments on the manuscript, and N. Buehner for technical advice. We are grateful to the NIH for grant R01-HD038921 to M.F.W, which supported this work.

## Author contributions

S.M. and M.F.W. designed the experiments; S.M. carried out the experiments; S.M. and M.F.W analyzed the results. S.M. and M.F.W. wrote and revised the manuscript.

## Conflict of interest statement

The authors declare no conflict of interests.

## Supplementary figures

**Fig S1.**
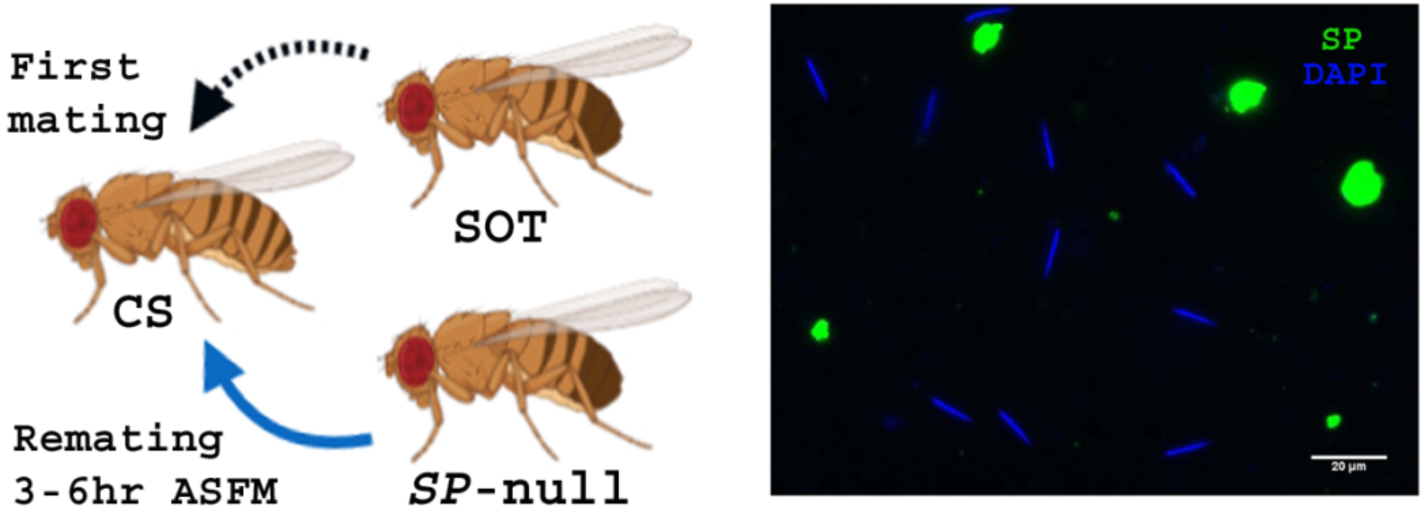
**Cartoon:** Pictorial representation of cross (fly images from Biorender). Females mated first with spermless (*sot*) male and then 3-6 hr ASFM with *SP*-null male that provided sperm. **Panel**: Sperm from SR of females mated to spermless males and then remated to *SP*-null males, 3-6hrs ASFM, frozen at 2hrs ASSM. Sperm heads were stained with DAPI (blue) and SP (green) probed with Alexa fluor 488 (n=5; Bar=20µm).

**Fig S2:**
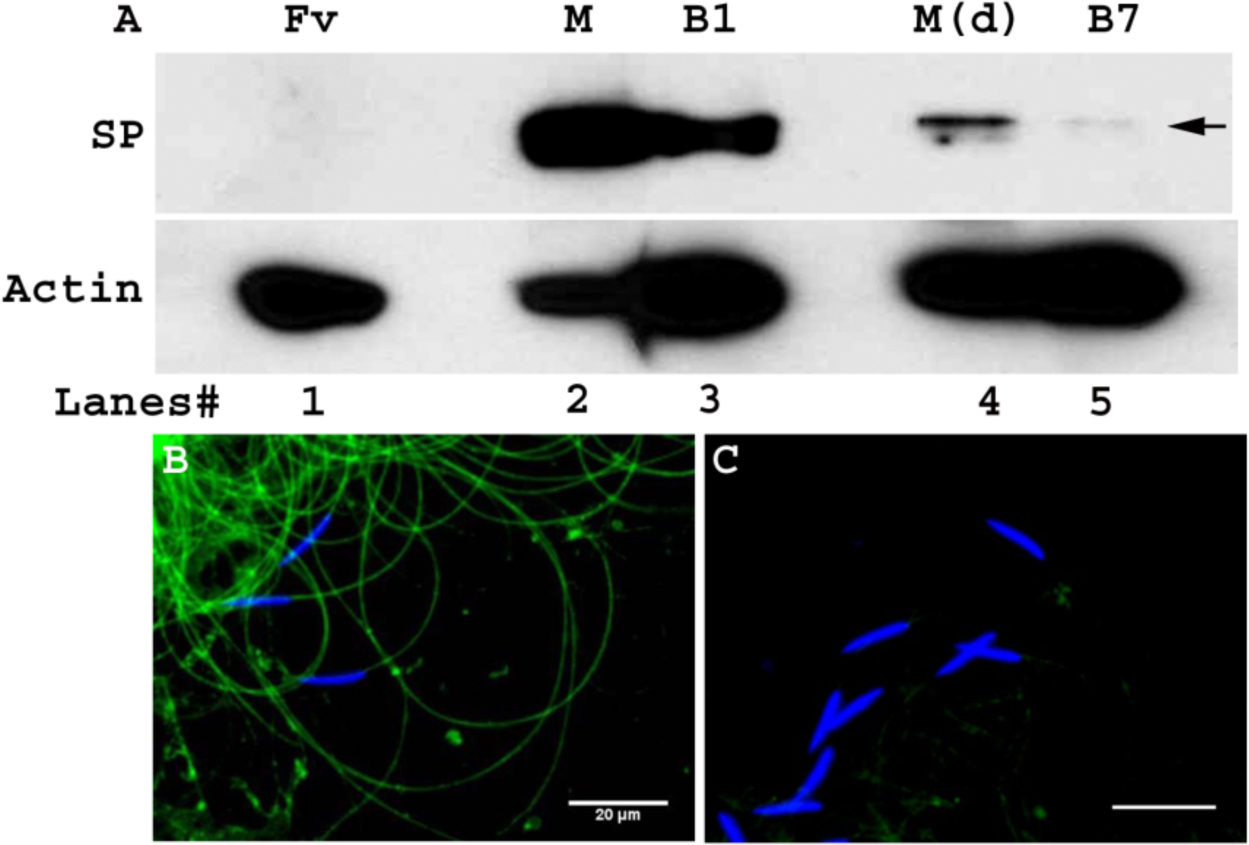
**(A)** Western blot probed for SP. Lanes/samples are, **1:** Fv, reproductive tract (RT) of 2 virgin females (negative control), **2:** M, 1 pair of male accessory glands from a 3 day old unmated virgin male, **3:** B1, RT of 4 females mated to control unmated virgin males, frozen at 2hr ASM, **4:** M(d), 1 pair of male accessory glands dissected from a multiply mated male (previously mated with six virgin females), **5:** B7, RT of 4 females mated to multiply-mated males, frozen at 2hr ASM. Actin served as loading control. **(B)** Sperm dissected from females mated to unmated males, frozen at 2hrs ASM. **(C)** Sperm dissected from females mated to multiply mated males, frozen at 2hrs ASM. Sperm heads were stained with DAPI (blue) and presence of SP (green) detected with Alexa fluor 488 (n=5; Bar=20µm).

**Fig S3:**
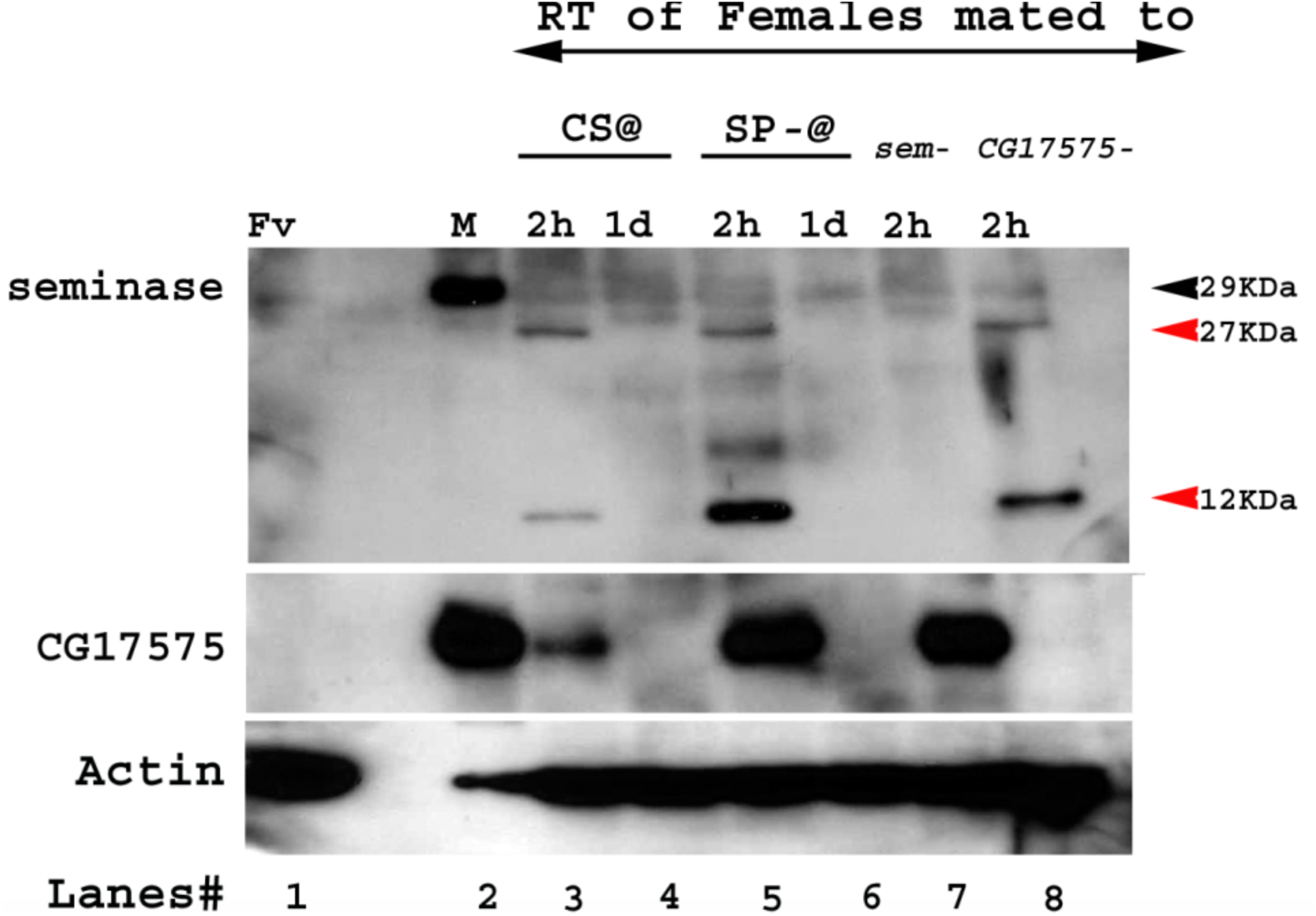
Western blot probed for seminase and CG17575. Lanes/samples are, **1:** Fv, reproductive tract (RT) of 3 virgin females (negative control), **2:** M, 1 pair of male accessory glands (positive control), **3-4:** RT of 5 females mated to wild type (CS) males at 2hr and 1d ASM, respectively, **5-6:** RT of 5 females mated to *SP-null; ProtB-eGFP* males at 2hr and 1d ASM, respectively **7:** RT of 5 females mated to *seminase*-null males at 2hr ASM **8:** RT of 5 females mated to *CG17575-*null males at 2hr ASM. Black arrows indicate full length seminase, red arrows indicate the cleavage products of seminase, post-mating in the female RT. Actin served as loading control.

**Fig S4:**
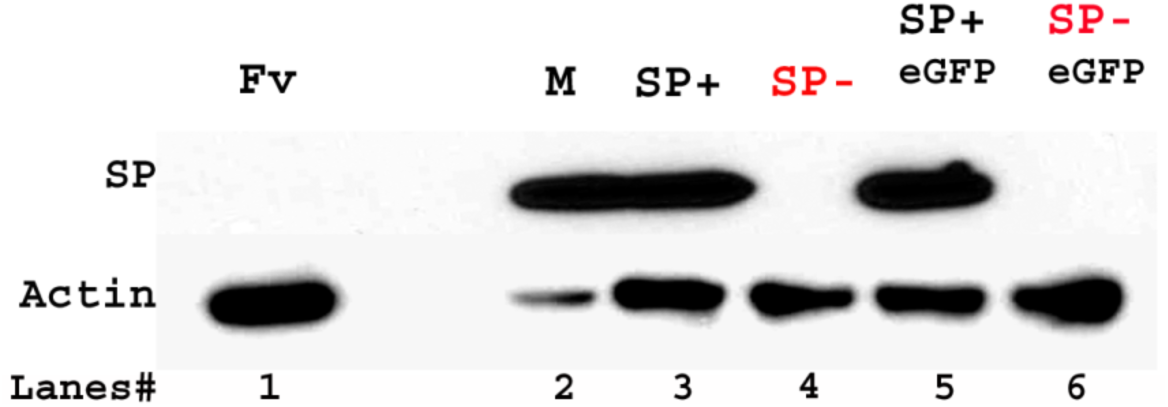
Western blot probed for SP. Lanes/samples are, **1:** Fv, reproductive tract (RT) of 3 virgin females (negative control), **2:** M, 1 pair of male accessory glands (positive control), **3:** SP+, RT of 3 females mated to control (TM3 siblings of *SP*-null males; positive control) males at 2hr ASM, **4:** SP-, RT of 3 females mated to *SP*-null males at 2hr ASM, **5:** SP+ eGFP, RT of 3 females mated to control (TM3 siblings of *SP-null; ProtB-eGFP* males; positive control) males at 2hr ASM **6:** SP-eGFP, RT of 3 females mated to *SP-null; ProtB-eGFP* males at 2hr ASM. Actin served as loading control.

## Notes

### Competing Interest Statement

The authors have declared no competing interest.

### Summary of Updates

some additional references and some clarifications in the text, and in the style of Fig. 3

